# BIOPRINTING OF MICRODISSECTED TUMOR “CUBOIDS” IN HYDROGELS

**DOI:** 10.1101/2025.09.05.671932

**Authors:** Anjul M. Bansal, Lisa F. Horowitz, Marcus Yeung, Taranjit S. Gujral, Albert Folch

**Affiliations:** Department of Bioengineering, University of Washington, Seattle, USA; Human Biology Division, Fred Hutchinson Cancer Research Center, Seattle, USA

**Keywords:** Bioprinting, Tumoroids, Hydrogels, Cancer

## Abstract

Microscale tumor models made from microdissected tumors that retain much of the original human tumor microenvironment (TME) are emerging as an alternative to preclinical animal models. We have introduced a drug testing approach that utilizes regularly-cut, cuboidal-shaped microdissected tissues, or “cuboids,” as a way to maximize creation of microtissues from scarce biopsy materials. However, microtissues (e.g., cuboids, organoids, spheroids, etc.) can be difficult to place in precise locations, especially in applications that require their culture in hydrogels. Here, using cuboids from mouse tumor models, we demonstrate a simple bioprinting strategy for precise placement and immobilization of cuboids in hydrogel. We use a commercial bioprinter to bioprint-containing hydrogel into arrays of small hydrogel dots containing cuboids, or “cuboid dots,” either onto a Transwell insert or into traps on a microplate. The hydrogel serves to immobilize the cuboids in place and provides a matrix to support cuboid viability. We demonstrate proof-of-concept applications for cancer drug testing and for protein profiling analysis. This approach will enable interface of cuboids with other devices, such as on top of a sensor or in a microfluidic platform. Furthermore, this automated process of dispensing and localizing cuboids (or other microtissue formats such as spheroids or organoids) could further their application to drug discovery and personalized medicine.

**TOC Graphic:** 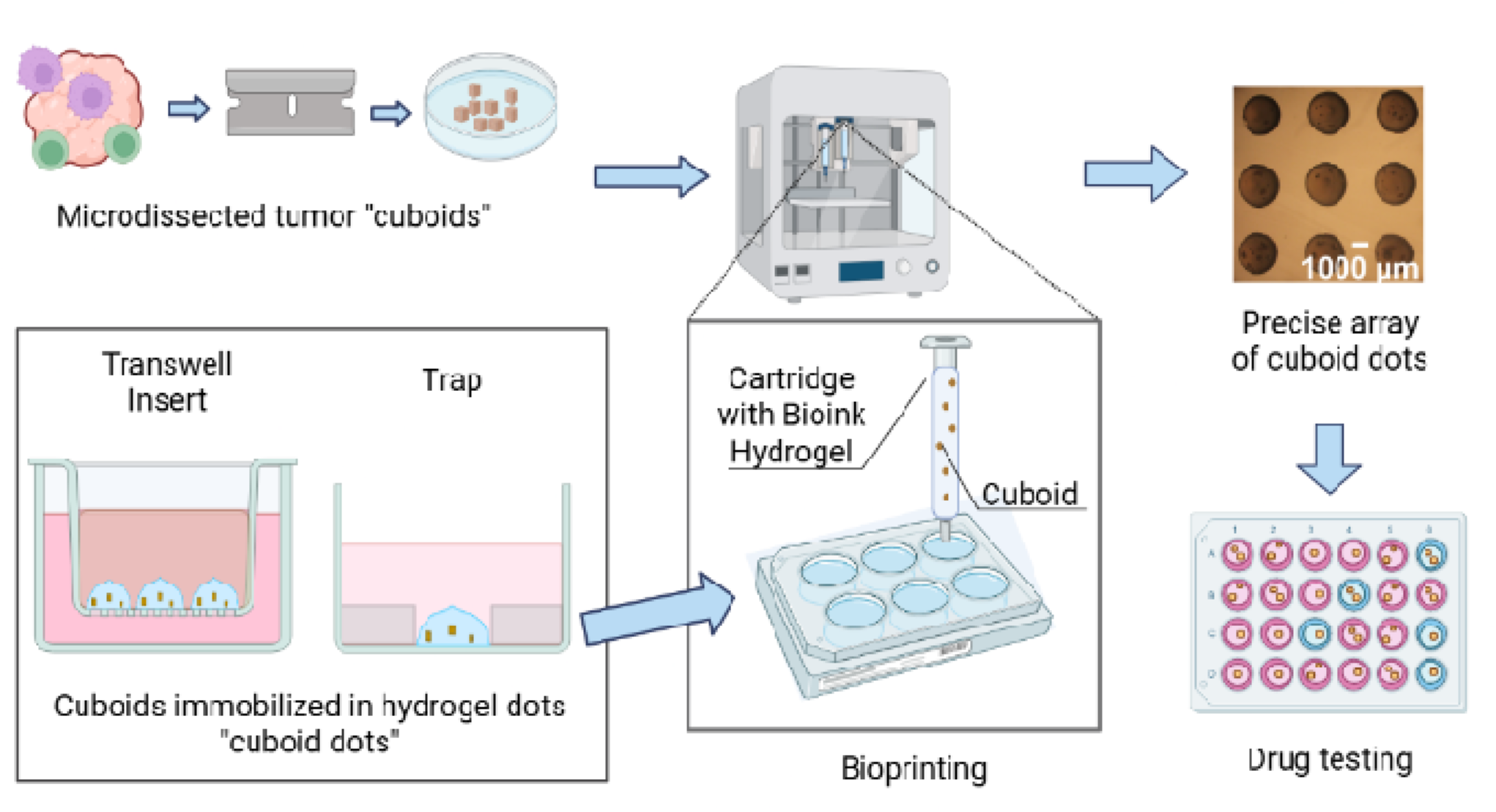

## 1. Introduction

Unlike animal models or tissue cell lines, microscale tumor models made directly from patient tumor biopsies can retain much of the complex and unique human tumor microenvironment (TME) and thus promise to serve as higher-fidelity drug tests for human cancers^1,2^. The most widespread and successful microscale tumor model, tumor spheroids (small spheres or “tumoroids”) created by dissociation of patient cells followed by expansion in culture, have cell-cell 3-D interactions that more closely resemble *in vivo* interactions than 2D cultures do^3–10^. Spheroids have been used for high-throughput drug screening assays that are predictive of some of the patients’ responses^2,3,10^. However, spheroids lose the original TME over time during amplification in culture, so drug responses that depend on the TME, such as immunotherapies, cannot easily be replicated in tumor spheroids.

In contrast, microdissected tumors – derived from cutting tumors into small pieces or slices – have shown promise as a microscale tumor model and drug testing approach that preserves the TME^2,11–16^. Some work has integrated microdissected tumors with microfluidics. Gervais’ group prepared cylindrical microdissected tissues and tested their response to cancer drugs with a microfluidic device^11^. The Barbie/Kamm group embedded minced tumor fragments in collagen within a microfluidic device. The Kuo group also showed response to checkpoint inhibitors, but with an air-liquid interface and collagen hydrogel in a Transwell instead of a microfluidic device^17^. Except for Gervais and our group, the majority of previous work on microdissected tumors has used slices or manually minced tissue fragments^12–14^; the Tang group has recently developed a micromachined grid to precisely cut tissues^18^. However, the random seeding of microtissues within relatively large volumes of hydrogel in these promising studies limits their ability to scale up to large numbers of drug conditions or to adapt to custom lab-on-a-chip applications.

Hydrogels – porous, crosslinked polymer chains that absorb relatively high amounts of fluid – have been extensively used in 3D culture systems, including tumor models^15^. Hydrogels help to mimic the *in vivo* tissue stroma and are favorable for cell encapsulation and expansion to enable tissue regeneration and cancer therapy^19^. Hydrogels can serve as a matrix to support and maintain biological functions, such as with immune cells, cancer tumor tissue, and various stem cells^20^. 3D hydrogel models can develop gradients, interrogate 3-D cell-cell and cell-ECM interactions, and mimic the microenvironment of a tissue, thereby offering more physiologically relevant models compared to 2D cell culture. These features make hydrogels an attractive matrix to support cuboids.

Bioprinters have been used extensively to bioengineer tissues by extruding cells in hydrogels^21^. Traditional uses of such extrusion bioprinting include building a complex biological structure with hydrogel, such as blood vessels, skin tissue, or organ-on-a-chip, with the main goal of recapitulating a functional biological microenvironment^22,23^. In numerous studies, extrusion of cells by a bioprinter creates organoids in hydrogel^24–26^, including tumoroids^24–27^. Volumetric printing of preformed liver organoids in hydrogel by light exposure without layers was shown to create liver-like structures^26^. Heinrich *et al*. incorporated glioblastoma cells and associated macrophages in hydrogel to develop a 3D bioprinted mini-brain model to study cellular interactions^27^.

Several bioprinting approaches have addressed precise placement of microtissues such as preformed (grown) spheroids. The Guo group used acoustic droplet bioprinting to precisely place spheroids in arrays of dots^28^. Although droplet-based bioprinting offers high precision and is nozzle-free, the droplets are limited in size to 10-150 µm dots, and cannot directly print spheroids^28,29^. To place larger spheroids up to 800 µm, the Ozbolat group used “aspiration-assisted” bioprinting (i.e., using pipettes as suction cups to lift individual, grown spheroids and place them into precise positions) within a continuous hydrogel bath with minimal loss of viability^30,31^. To directly print spheroids within a hydrogel, the Swaminathan group used extrusion-based bioprinting to print 100 µm breast cancer (grown) spheroids in various hydrogels, validating their viability and response to drugs^32^. However, due to nozzle clogging, extrusion-based bioprinting has been limited to spheroids of less than 400 µm^32,33^. To the best of our knowledge, no studies have demonstrated direct extrusion bioprinting of large (∼400 µm) spheroids or intact microtissues in hydrogel.

Here we performed extrusion bioprinting of mouse breast cancer cuboids (i.e., cuboidal-shaped microtissues with an intact TME) encapsulated in hydrogel dots, or “cuboid dots,” in different array patterns. With this process, we immobilized mouse tumor cuboids in defined spatial positions on Transwell membranes as well as in custom microtraps in a microdevice. We then demonstrated the use of cuboid dots for drug testing with a cell death assay as well as with Reverse Phase Protein Profiling (RPPA) which analyzed molecular responses to drugs within cuboids. To the best of our knowledge, this work represents the first effort in bioprinting hydrogel-embedded intact-TME microtumors. In the future, such “cuboid dots” could provide a simple means to interface microtumor models with more complex devices, such as a tumor-on-a-chip model or a cancer biosensor for personalized oncology.

## 2. Experimental (Materials and Methods)

### 2.1. Materials

We purchased the hydrogels Startink, Cellink, and GelMA, and the crosslinker CaCl_2_ from CellInk (Gothenburg, Sweden). Cell lines to generate mouse syngeneic breast cancer tumors (PY8119) or human glioma xenograft tumors (U87-MG) were purchased from ATCC (Manassas, Virginia.) We obtained C57BL/6J mice (syngeneic models) and *Foxn1^nu^*athymic mice (xenograft model) from Jackson Laboratories (Bar Harbor, Maine). For PMMA plastics we utilized 0.8 mm-thick sheet from Astra Bioproducts (Copiague, New York) and 6.35 mm-thick sheet from McMaster-Carr (Elmhurst, Illinois).

### 2.2. Bioprinting materials

We used CellInk’s Inkredible+ bioprinter and three of CellInk’s bioprinting hydrogels. CellInk’s Bioink, a semitranslucent, insoluble gel (viscosity 7000 Pa·s) made of alginate and cellulose fibers, maintains its shape and stability in solution after crosslinking with CaCl_2_. CellInk’s Start, a transparent, water-soluble gel (polyethylene oxide, viscosity of 157 Pa·s) intended for use as a sacrificial material, dissolves easily in liquid such as media. CellInk’s GelMA, a bioink made of a mixture of porcine gelatin functionalized with methacrylate groups and photoinitiator, maintains its shape in solution after being photocrosslinked (Table 1).

**Table 1.**
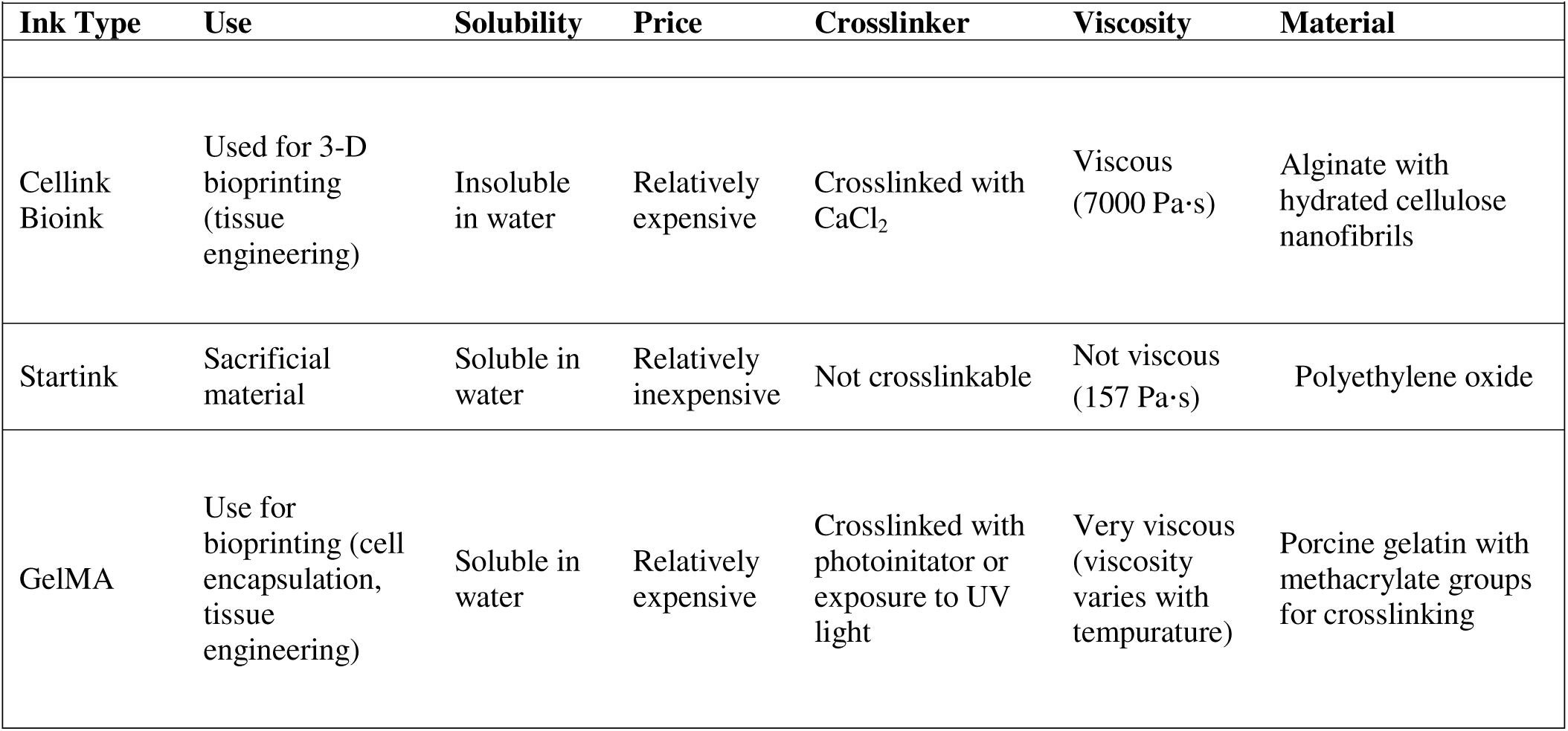
Types of Bioink used to print (CellInk)

### 2.3. Cuboid generation and culture

All mouse experiments were performed according to recommendations by the institutional guidelines at the University of Washington or at the Fred Hutchinson Cancer Research Center. We prepared ∼400 µm-wide cuboids from mouse syngeneic breast cancer tumors (Py8119, ATCC) or from human glioma xenograft tumors (U87-MG, ATCC) as described in our previous work^34^. Briefly, cell lines were grown in DMEM/F12 (Invitrogen) supplemented with 5% fetal bovine serum (VWR) and penicillin/streptomycin (Invitrogen) and passaged twice a week. We injected female mice aged 6-10 weeks-old subcutaneously in the flank with 1-2 million cells in 200 µL of DMEM-F12 medium (U87) or into the mammary fat pad with 1-2 million cells in 50 µL of Matrigel (Corning) (Py8119). We sacrificed mice before the tumor volume reached 2 cm^3^.

To generate microdissected cuboids, we first glued small pieces (<0.5 cm) of the tumor biopsy onto a PMMA disk with cyanoacrylate glue in a thin layer below as well as around the edges of the biopsy. These thin layers of cyanoacylate glue have no effect on cuboid viability. The tumor was cut into slices, then into cubes (∼400 μm × 400 μm × 400 μm) with a tissue chopper device (McIlwain Tissue Chopper, Ted Pella Inc.)^34^. Alternatively we did the first step to create slices using a MZ5100 vibratome (Lafayette Instruments) after embedding tissue punches (6 mm diameter, Harris Uni-Core) in 1-2% low-melt agarose. To select the size of the cuboids, we passed them first through 750 μm-size mesh filters and then through 300 μm-size ones (Pluriselect). We cultured the cuboids in growth medium (DMEM/F12, 5% heat-inactivated FBS, 1% penicillin/streptomycin, Invitrogen) for up to 3 days before bioprinting.

### 2.4. Bioprinting procedure

We prepared cuboids in hydrogel (either 250 or 500 cuboids/mL of bioink) as follows. First, we mixed a volume of bioink (ranging from 0.75 mL to 1.5 mL) in a petri dish with the appropriate number of cuboids (in 10 µL of either DMEM/F12 media or PBS). We then transferred the mix into a 3 mL syringe and loaded it into a cartridge (CellInk) using a female-female Luer lock (Quosmedix).

To enable direct locational control of the bioprinter, we wrote files in G-code using the CellInk Heartware (Inkredible+’s operating software). We placed dots 6 mm apart and across for the 3 × 3 Transwell array and 9 mm apart for the 6 × 4 microplate array. The bioprinter is reported to have a precision of up to 10 µm. We corrected for a misalignment of the axes of the bioprinter that caused a slight downwards vertical shift while printing horizontally. We empirically determined the equation for the correction as Y(X) = -0.00604 · X + Y(0).

We printed arrays of hydrogel cuboid dots using an Inkredible+ bioprinter from CellInk (Gothenburg, Sweden). We used a 14-gauge pin (Loctite) as a nozzle with 1.6 mm inner diameter and a temperature of 25 °C. For larger-sized dots, we set the extrusion pressure to 20 kPa and printed for 1.5 sec. For smaller sized dots, we set the pressure to 13 kPa and printed for 2 sec. When printing larger numbers of dots at once (24 cuboid dots for the microdevice), we observed progressively larger cuboid dots (48% increase in volume per 30 sec, ∼2 sec per dot). To compensate, we manually turned down the extrusion pressure by 1 kPa for the last row of printing. For the microwell printing, we printed the smaller dots on a 6 × 4 array in a custom microplate array with traps. At the end, we immersed the dots in 50 mM CaCl_2_ cross-linking solution (CellInk) for 10 min. For printing onto Transwell inserts, we printed the larger-sized dots on a 3 × 3 array on top of a Transwell insert (Millipore) in a 6-well plate. We then added the 50 mM CaCl_2_ solution to the bottom of the inserts to crosslink the cuboid dots from the bottom for 10 min. Regardless of the application, after the 10 min incubation with crosslinker, we rinsed the cuboid dots 3 times with PBS.

### 2.5. Microplate fabrication

We fabricated two variations of microplate devices with microtraps, one with and one without an additional 24-well layer, which was used for drug testing. We utilized CO_2_ laser micromachining (VLS3.60, Universal Laser Systems) and bonding as detailed in Horowitz *et al.*^34^. The microtrap layer consisted of 0.8 mm-thick sheet of PMMA (Astra Bioproducts) with 2 mm-radii traps. A border and an insertable bottomless 24-well plate layer, both 3.5 cm-wide by 5.5 cm-long, were made with 6.35 mm-thick PMMA (McMaster-Carr). We attached the bottom sealing layer by thermal-solvent bonding at 140 °C after exposure to chloroform vapor. We attached the border, and the well plate layer for experiments with drugs, by adhesive bonding with pressure-sensitive adhesive (300 LSE, 3M).

**Caution**! Toxic chloroform gas is a potential carcinogen. The entire apparatus was set-up in a fume hood. Standard lab personal protection equipment were worn when working on the pressurized system. The sash was always as low as practical for the work.

### 2.6. Cuboid viability and drug treatment assays

For the bioink viability assay, we performed a RealTime-Glo (Promega) assay according to manufacturer’s instructions. We manually extruded the ink containing cuboids with a 3 mL syringe and 14 gauge pin as a nozzle into 6-well plates. Luminescence was measured by IVIS (Perkin Elmer) at day 1 (baseline) and at day 3.

For drug treatments, we treated cuboids with drugs (2 µM staurosporine, 10 µM cisplatin, 100 µM cisplatin, 2 µM doxorubicin, or 30 µM mocetinostat (Medchem Express)) for 3 days with cuboid dots completely covered by solution. We treated the cuboid dots with 0.05% DMSO as vehicle control. To perform the cell death assay, we exposed the cuboid dots to fluorescent dead nuclear stains, SYTOX green (SG; Invitrogen, 0.01 µM) or Reddot2 (1:400 dilution, Invitrogen) for 2 hrs, followed by three washes with PBS. After overnight fixation of the cuboid dots in 4% paraformaldehyde and washing twice in PBS, we captured brightfield and fluorescent images at 2x magnification with an epifluorescence microscope (BZ-X810, Keyence). We quantified the fluorescence of each cuboid using Fiji^35^.

We performed a reverse phase protein array (RPPA) on Py8119 cuboids after drug treatment. RPPA is a method that reveals the activation state of different protein pathways using panels of antibodies against phosphorylated proteins, quantifying their value. We used cuboids physically removed from cuboid dots after treatment, cuboids still in the cuboid dots, and non-bioprinted cuboids, all cultured in parallel. We froze (-80 °C) approximately 15 cuboids for each condition in 1.5 mL tubes. In brief, we prepared lysates from each sample onto a 2% SDS lysis buffer^36^. We printed these out onto RPPA slides, each containing 16 nitrocellulose pads (Grace Biolabs, GBL505116). We fitted these slides to hybridization chamber cassettes (Gel Company, AHF16) and incubated them with primary and secondary antibodies. These samples were then scanned using Licor Odyssey CLx Scanner (LiCOR), and the total signal intensity was quantified using Array-Pro analyzer software package (Media Cybernetics, Maryland, USA). The results from the control cuboids (one antibody) were included in supplemental data for another publication^37^.

### 2.7. Statistical analysis

Data analyses were performed using Graphpad Prism version 9.5. Descriptive statistics, including means and standard deviations, were calculated to summarize the sample characteristics. A one-way analysis of variance (ANOVA) was conducted to assess differences among groups. A Dunnett’s multiple comparisons test was used to compare each treatment group to the control group to control for Type 1 error. Statistical significance was set at α = 0.05 for all analyses.

## 3. Results

## 3.1. Bioprinting of cuboids in hydrogel

We bioprinted cuboids in hydrogel with the Inkredible+ bioprinter, a pneumatic-based extrusion system with dual printheads that allows for bioprinting cell-laden hydrogels. To print the relatively large cuboids (∼400-µm diameter versus <30 µm for cells), we used large diameter pins (14 gauge, 1.6 mm inner diameter) instead of the provided tips (23-27 gauge, 0.4-0.2 mm ID). Additionally, to allow us to print in defined locations, we ran custom G-code routines instead of using preset code on the provided software. Our custom G-code routine allowed us to set the time and speed of ink extruded to get a fixed dot size. Furthermore, by setting our own distance between subsequent cuboid dots, we were able to create minimally spaced cuboid dots that do not touch.

We used three different approaches to bioprint arrays of hydrogel spots containing cuboids, or “cuboid dots,” at predetermined, arbitrary positions. An overview of these three methods is shown in **Fig. 1**. Each method corresponded to a different surface: directly on a well plate (**Fig. 1**A), on top of a Transwell insert (**Fig. 1**B), and onto laser-cut traps in a microwell device (**Fig. 1**C). When we printed the cuboid dots directly on top of the plate, changing the medium often caused the cuboid dots to detach and become displaced from their original position (**Fig. 1**A). When we printed the cuboid dots on top of a Transwell insert and crosslinked the hydrogel from underneath the Transwell (50 mM CaCl_2_ cross-linking solution for 10 min, see Methods), the cuboid dots remained attached and did not change position (**Fig. 1**B). We also printed smaller dots in a custom microplate with traps (0.8 mm-deep microwells) that immobilized the cuboid dots at defined positions (**Fig. 1**C). For all three methods, we cultured the cuboid dots by submerging them in media for ease of comparison. Note that the Transwell method, in principle, allows for cuboid culture with an air-liquid interface, with medium from below the membrane and air from above. Both the Transwell crosslinking method and the microplate localization method allowed for cuboid culture within a supportive hydrogel environment with localization to a regular array. On the other hand, the direct printing on a dish facilitated cuboid culture in hydrogel for applications that required cuboid dot retrieval for downstream analysis.

**Figure 1.**
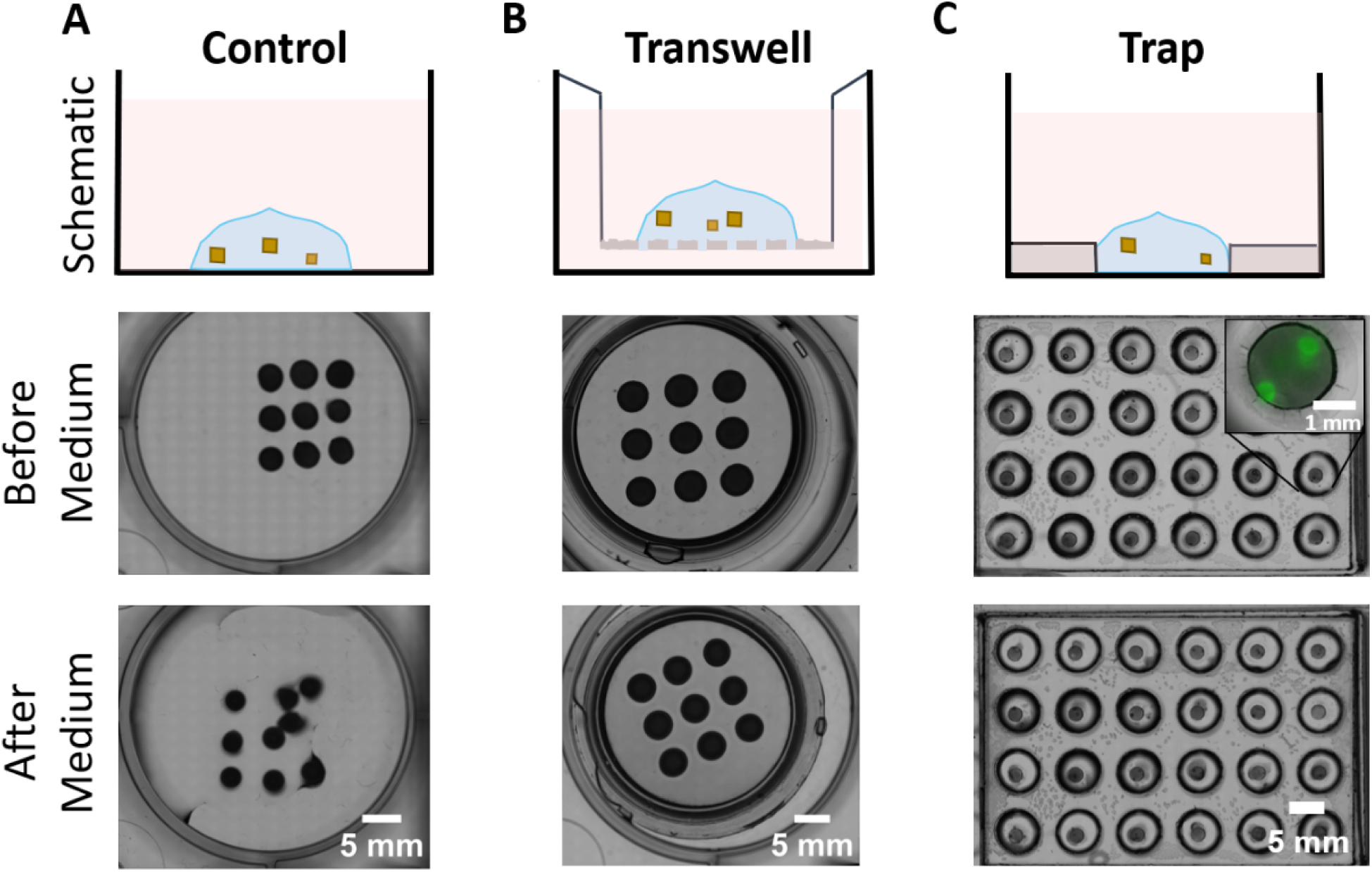
Three methods to bioprint cuboids in hy drogel cuboid dots: The top row depicts schematics of the final layouts of the cuboids (brown) in hydrogel (blue) submerged in medium (pink). The medium and bottom row show various conditions before and after incubation in cell culture medium. (A) A 3 × 3 array of cuboid dots directly seeded onto a tissue culture well plate; note how the cuboid dots did not maintain their position once in medium (bottom image). (B) A 3 × 3 array of cuboids seeded on top of a Transwell insert; note how the dots maintained their position after the addition of medium (bottom image). (C) A 6 × 4 array of hydrogel cuboid dots printed into circular traps in a microplate, with one trap per well; note how the dots maintained their position after the addition of medium (bottom image). A fluorescent overlay (inset, middle image) highlights that cuboid dots in a specific trap can

### 3.2. Bioprinting parameters for cuboid dots

Changing different bioprinting parameters allowed us to control the size of the dots (**Fig. 2**A). By optimizing printer extrusion time (1, 1.5, and 2 seconds), and pressure (16 and 18 kPa), we created hydrogel dots (Cellink Bioink) with sizes ranging from 0 mm to 2 mm radii (**Fig. 2**A). We measured volume (gel density 1 g/mL) by weighing nine dots per condition. The dot volume did not increase linearly with dot radius. After plotting extruded hydrogel dot volume V(*r*) as a function of dot radius *r*, we fit the curve V(*r*) =2.193*r*^2.452^, with an R^2^ value of 0.99 (**Fig. 2**B). We also observed that the hydrogel dots swelled with exposure to culture medium. The initial hydrogel dot volume was 9.3 ± 1.6 µL, (ave ± s.d.; measured by weight), corresponding to a *r* =1.8 mm calculated from the curve fit (Ca^2+^ crosslinked dots, n=6 sets of 3 hydrogel dots each). The dots swelled in PBS solution to about 4.3 times its size (40 ± 7 µL, n=6 sets of 3 hydrogel dots) after one hour and ∼11 times its size after 3 days (n=3 sets of 3 hydrogel dots).

**Figure 2.**
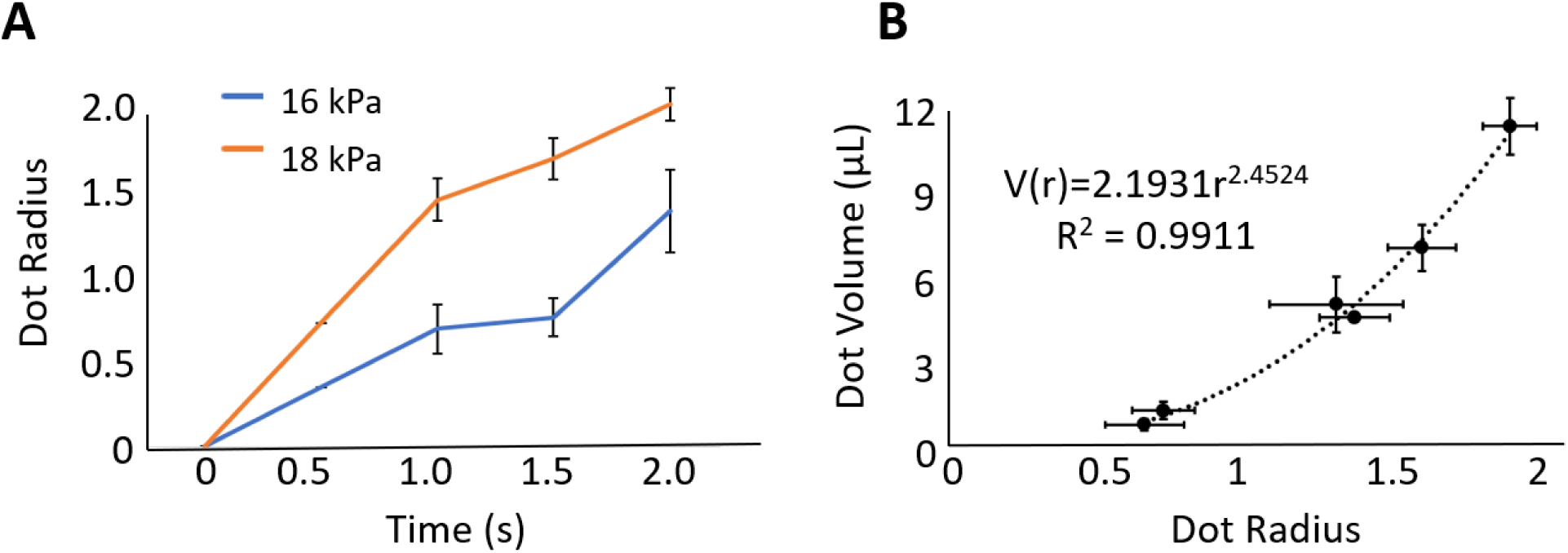
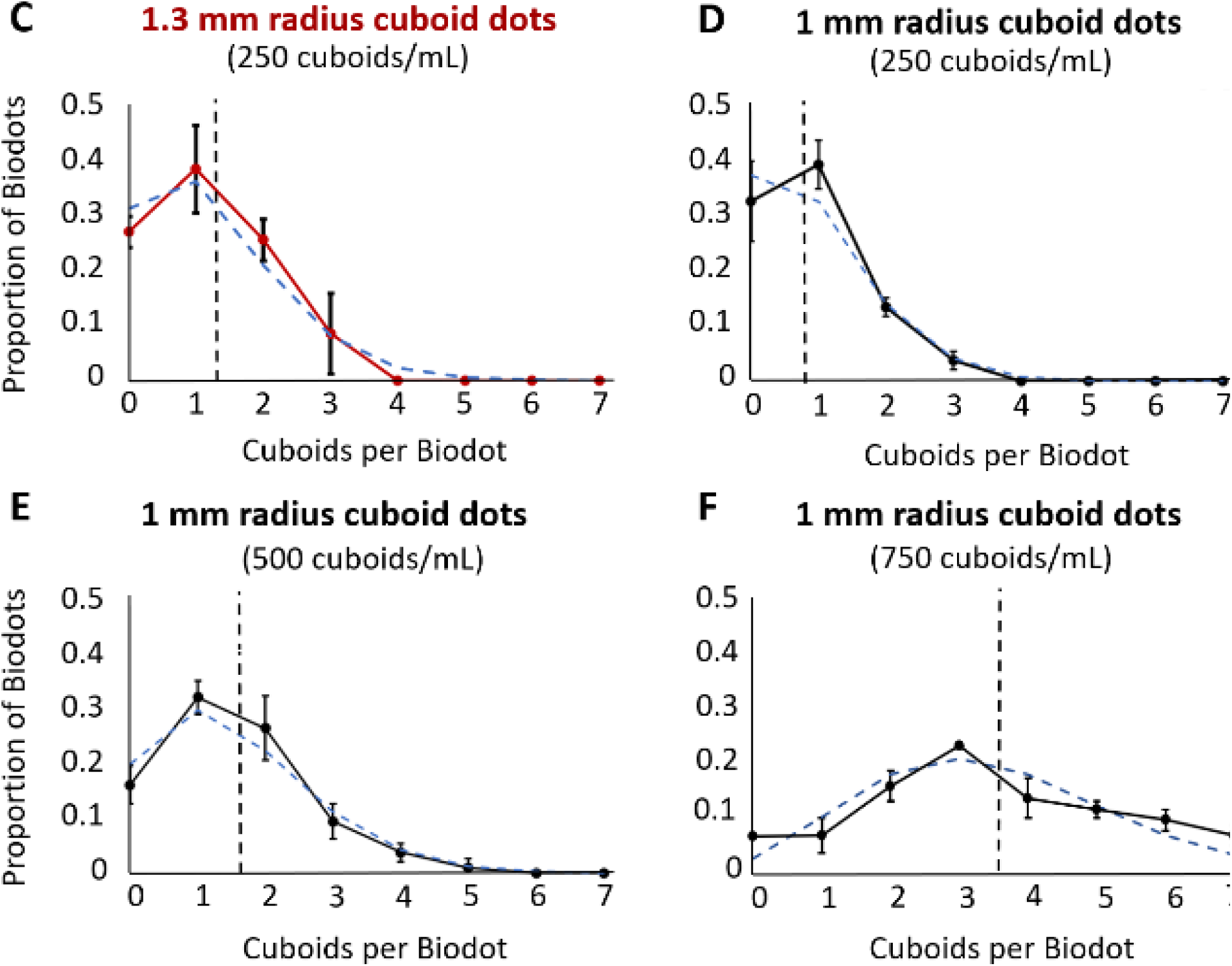
Control of size and cuboid distributions for cuboid dots: (A) Cellink dot radius as a function of extrusion time shown for two different pressures, 16 kPa and 18 kPa (n=9 per condition). (B) Dispensed volume as a function of hydrogel dot radius (n=8-9). (C-F) Histograms showing the distribution of cuboids per cuboid dot for different cuboid concentrations and dot sizes. In each graph, the black vertical dashed line denotes the average number of cuboids per dot and the superimposed Poisson distribution with λ=average is shown in blue dashed line. (C) 1.3 mm radius cuboid dots using a hydrogel solution at a concentration of 250 cuboids/mL. Average number of cuboids per dot = 1.2 ± 0.9 (n=70). (D) 1 mm radius cuboid dots using a hydrogel solution at a concentration of 250 cuboids/mL. Average number of cuboids per dot = 0.87 ± 0.82 (n=93). (E) 1 mm radius cuboid dots using a hydrogel solution at

We then evaluated our ability to control the number of cuboids per dot using Cellink Bioink (**Fig. 2**C-F). First, we printed larger cuboid dots with ∼1 cuboid per dot on Transwell inserts in a 6-well plate, which has more space for larger cuboid dots as well as for media or drugs (**Fig. 2**C). To print an average of 1 cuboid per 1.3 mm hydrogel dot we used a concentration of 250 cuboids/mL, or 1 cuboid/4 µL (volume measured by weight: 4 ± 0.97 µL; ave ± s.d., n=3 with 9 dots weighed for each sample). With these conditions, on average each bioprinted dot had 1.2 ± 0.9 cuboids (n=70); 39% had 1 cuboid, 26% had 2 cuboids and 27% had no cuboids. Next, we varied the cuboid number in 1 mm-sized cuboid dots (volume approximated by curve fit, V(*r*) = 2.2 µL) printed inside the small wells of a microplate (96-well plate well dimensions, 9-mm diam.) (**Fig. 2**D). To print mostly one or no cuboids per dot, we used a cuboid concentration of 250 cuboids/mL, which represents an expected 0.55 cuboid/dot. Using these parameters, 44% had 1 cuboid, 16% had 2 cuboids, and 37% had no cuboids, resulting in an average of 0.87 ± 0.82 cuboids per dot (n=93). To print mostly 1-2 cuboids per 1 mm cuboid dot and minimize dots without cuboids, we used a concentration of 500 cuboids/mL (estimated 1.1 cuboids/dot; **Fig. 2**E). For this two-fold higher concentration, 66% had 1 or 2 cuboids, 18% had no cuboids, and 16% had 3 or more, resulting in an average of 1.5 ± 1.1 cuboids per dot (n=94). Lastly, to print multiple cuboids per dot, we used a concentration of 750 cuboids/mL (estimated 1.65 cuboids/dot; **Fig. 2**F). At this highest concentration, 85% had 2 or more cuboids, 7.5 % had 1 cuboids per dot, and only 7.5% had no cuboids, resulting in an average of 3.5 ± 1.1 cuboids per dot (n=72). Despite our limited ability to evenly distribute the cuboids within the very viscous hydrogel, the cuboid distribution using each method appeared to follow a Poisson distribution. Thus, by adjusting the cuboid concentration and/or dot size, one could further customize cuboid dots for particular purposes.

We also evaluated the biocompatibility of various hydrogel bioinks with cuboids. We performed a viability assay (Realtime Glo luminescence) with cuboids in bioinks extruded with a syringe compared to untouched control cuboids in medium alone (**Fig. S1**). After culture for 3 days, cuboids in Startink and in Cellink (with and without Ca^2+^ cross-linking) had similar luminescence ratios as the control, indicating similar viability. To confirm that the bioprinting extrusion process does not cause cuboid cell death, we compared the viability of bioprinted cuboids to non-printed cuboids in media (**Fig. S2**). These cuboids were from PY8119 mouse breast cancer tumors. We did not find evidence of cuboids being damaged by the extrusion process (n= 11-31 cuboids). Because identification of individual cuboid signal was too difficult in the viscous GelMA bioink, we compared the bulk luminescence signal and found that the overall viability was as good as or better than control (1.5 for GelMA, 0.9 for Cellink, 0.9 for control; post/pre). We did not further pursue GelMA for biodots because of its temperature-dependence of printing as well as its poor optics. We conclude that bioprinting of cuboids in all three of the hydrogel bioinks does not damage their viability. While Startink can serve for cuboid placement when the hydrogel should then be washed away, all subsequent experiments on cuboid placement within a stable hydrogel matrix were performed with Cellink.

### 3.3. Cuboid dot immobilization by microtraps

Microtrap wells cut into plastic represent an integrated method to immobilize cuboid dots without the need for expensive additional Transwell inserts. We optimized the microtrap dimensions for physical immobilization of Cellink hydrogel dots. We fabricated and tested different sizes of microtraps to find which size best prevented the detachment of Cellink dots during solution changes and movement of the plates, like those that occur during experiments with cuboid dots The 24-well microplate was designed for compatibility with drug testing, with a single microtrap at the center of each well in a 96-well plate format. To optimize the microtrap size, for these experiments without drugs we fabricated a test microplate without culture wells. The test microplate had an array of six 0.8 mm-high cylindrical microtraps of different radii (4 each, radii 0.125, 0.25, 0.5, 1, 1.5, and 2 mm). After printing, Cellink dots (without cuboids) with 1-mm radii in each of the wells, we tested the durability of their fixation. We subjected them to four stages of treatment: crosslinking with CaCl_2_, washing once with PBS, washing two more times with PBS, and shaking for 1 hr. After each stage, we counted the number of immobile dots for each trap size (**Fig. 3**A). The 1-mm radii traps best preserved the cuboid dots’ location as they held 100% of the cuboid dots in place throughout (**Fig. 3**B). As 1 mm is the size closest to the dot size, we hypothesize that the 1-mm traps gripped the dots with the most surface area along the vertical walls as compared to the other sizes. Traps with larger (1.5- and 2-mm radii) failed to grip the dots, making them lose position with just a change of solution. Traps that with smaller radii (0.125, 0.25 and 0.5 mm) fixed the dots’ position to some extent, but failed to withstand larger forces, such as those encountered during shaking. Hence, for optimum immobilization of ∼1 mm hydrogel dots during subsequent cuboid drug experiments, we fabricated our microplate with 1 mm-radii uniform traps centered at the bottom of each well.

**Figure 3.**
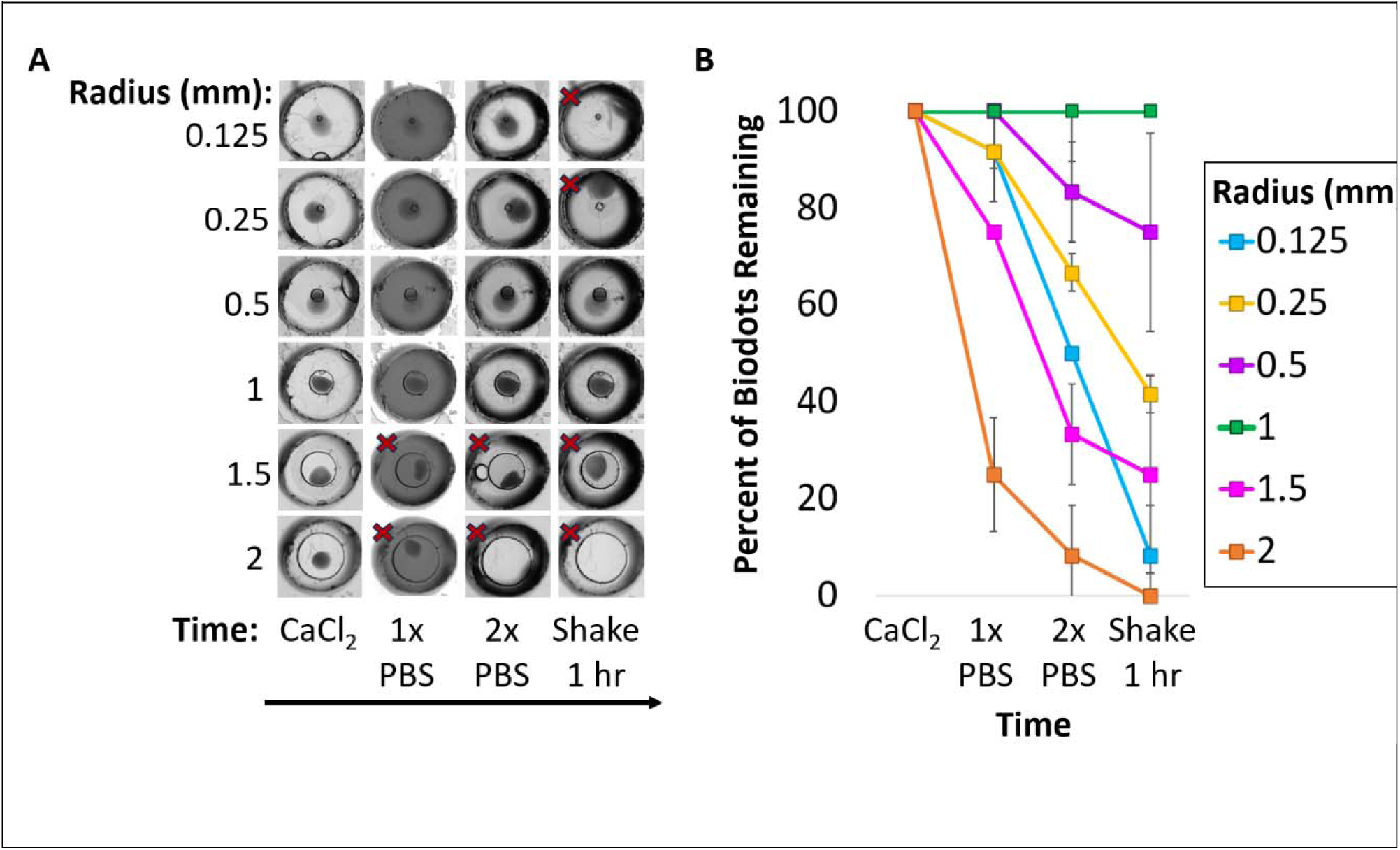
Trap Size Determination: (A) Individual examples of 1 mm-radius hydrogel dots printed on a microplate with differing trap widths (depth 0.8 mm) passed through 4 steps of treatment. Red ×’s indicate the dots that moved. (B) Graph of the percentage of dots remaining at each stage. The average dot radius is 1 ± 0.6 mm (n=12 per condition).

### 3.4. Cuboid drug treatment on Transwell insert

Next, we tested cuboid drug response on arrays of bioprinted cuboid dots that had been crosslinked to a Transwell insert (**Fig. 4**). Each well was printed with an array of 9 cuboid dots on a Transwell insert and received one drug treatment. We treated the Py8119 breast cancer cuboid dots with two cytotoxic chemotherapy drugs for three days: cisplatin at 10 µM and 100 µM (**Fig. 4**A) and doxorubicin at 2 µM (**Fig. 4**B). Cuboids treated with two different concentrations of cisplatin had higher levels of the SYTOX Green fluorescent cell death indicator than the control, indicating cell death (**Fig. 4**C). Similarly, cuboids treated with doxorubicin had higher levels of the Reddot2 far red cell death indicator (to not overlap with doxorubicin’s own fluorescence) than the DMSO vehicle control, indicating cell death (**Fig. 4**D). Thus, cuboids in bioprinted cuboid dots on Transwells showed clear drug responses by a fluorescence cell death assay.

**Figure 4.**
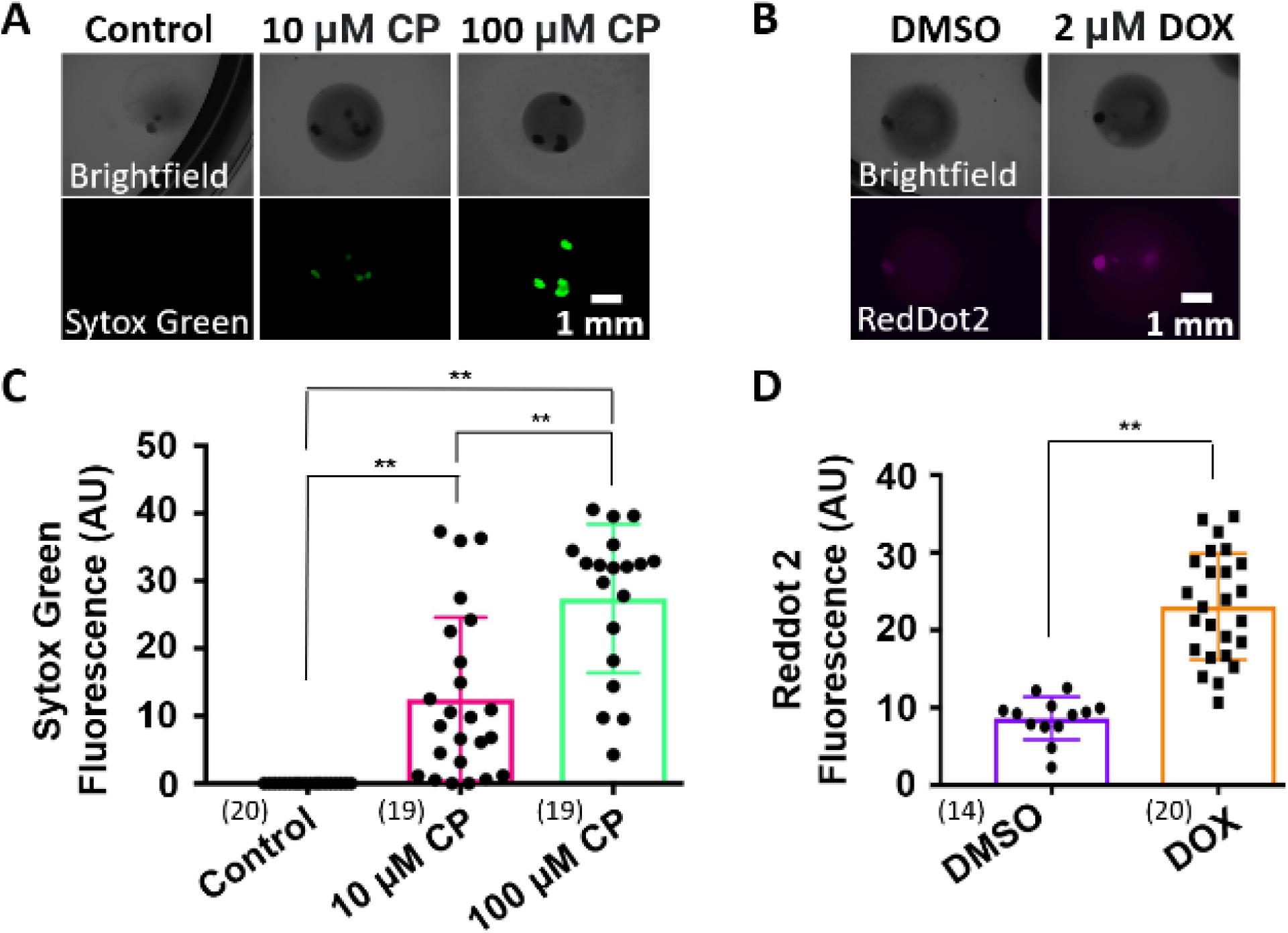
Drug treatment on Transwell inserts: (A) Brightfield and Sytox Green nuclear cell death dye images of Py8119 tumor cuboids in cuboid dots after 3 days of treatment with cisplatin or medium control. (B) Brightfield and RedDot2 nuclear cell death images after 3 days of treatment with doxorubicin or DMSO vehicle control. (C) Graph of cell death after cisplatin treatment. (D) Graph of cell death after doxorubicin treatment. Number of cuboids per condition shown in parentheses underneath each condition. * = p<.05 **=p< .01 One-way ANOVA with Dunnett’s Multiple Comparison test.

### 3.5. Cuboid drug treatment in a microplate

To demonstrate drug testing at the level of individual cuboid dots, we used a 24-well microplate with one microtrap and cuboid dot per well (**Fig. 5**A). We exposed cuboid dots containing U87 cuboids for 3 days to drugs: cisplatin (CP, two concentrations: 10 and 100 µM), mocetinostat (MOC, chemotherapy drug, histone deacetylase inhibitor; 30 µM concentration), staurosporine (STS, non-specific kinase inhibitor and apoptotic cell death drug; 2 µM concentration), DMSO vehicle control (for MOC and STS), or medium control (for CP) (**Fig. 5**B,C). Drug treatment led to increases in cell death, as seen by an increase in SYTOX green dead nuclear fluorescence (**Fig. 5**C). We note that cuboid dots increased in size over time (volumes of ∼4.2 µL initially and ∼43 µL after 3 days estimated from our previous swelling calculations). While the effect on drug concentration was minimal with immediate application as done here (decrease of 3% with 120 µL drug volume and ∼4.2 µL dot volume), treatments applied later would need to take into account the drug dilution by the liquid in the hydrogel dot. For the cell death dyes added at the end of the experiment, the final concentration was still sufficiently high despite the 33% decrease. In conclusion, we show that the cuboid dots in a microplate offered a more compact and scalable approach to test multiple drugs.

**Figure 5.**
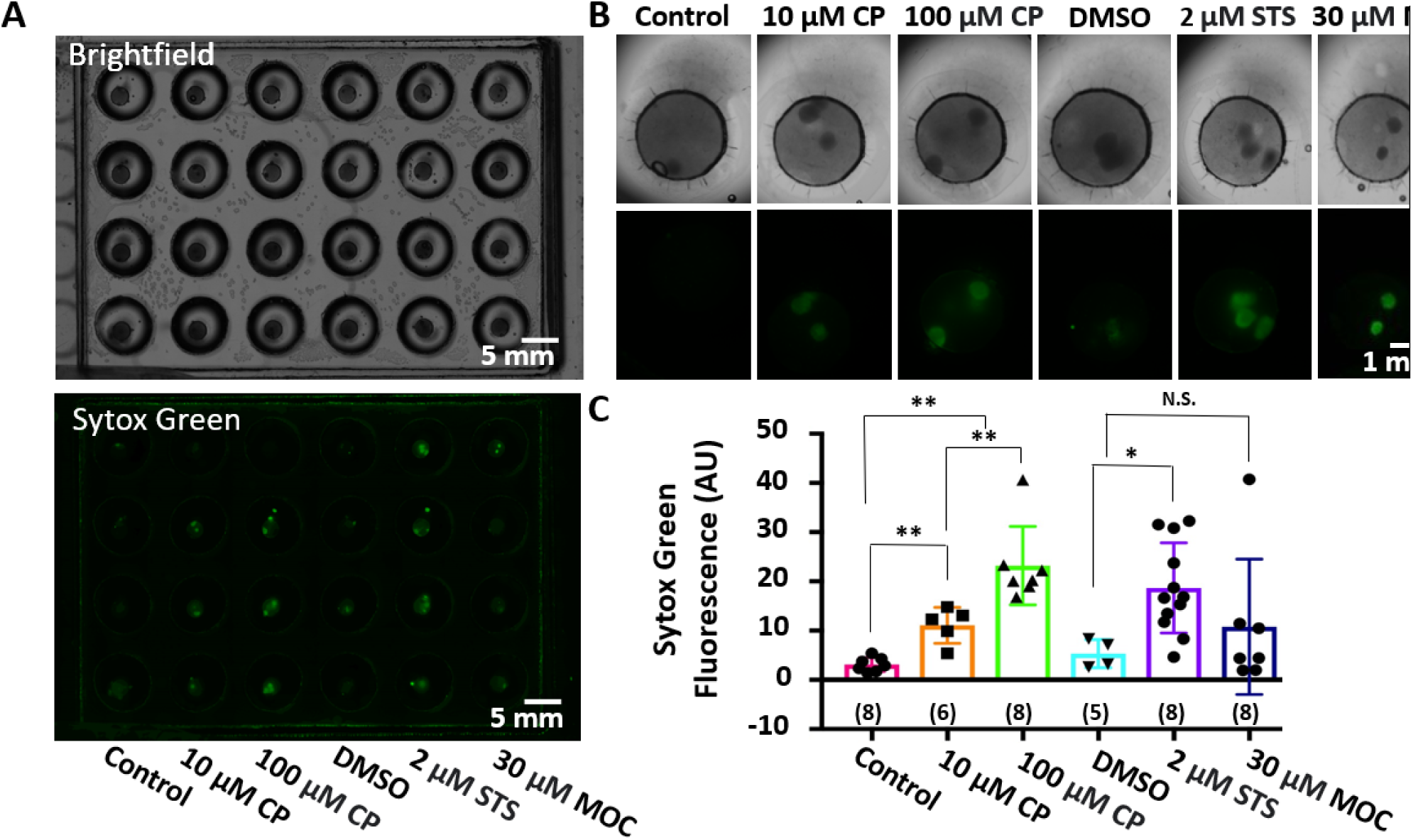
Cell death assay using microtraps: (A) Brightfield and Sytox green (green nuclear cell death) images of U87 cuboids on a microplate after treatment for 3 days with drugs: cisplatin (CP) compared to media control, staurosporine (STS) and mocetinostat (MOC) compared to DMSO vehicle. (B) Closeup of wells shown in (A). (C) Graph of fluorescence levels of cuboids treated with drugs. Number of cuboids per condition shown in parentheses underneath each condition. Average ± SEM *p < 0.05 **p < 0.01. One-way ANOVA with Dunnett’s Multiple Comparison test.

### 3.6. Molecular changes demonstrated on cuboid dots

To demonstrate a deeper analysis of the molecular responses to drugs within cuboids in cuboid dots, we assessed changes in phosphoprotein in the cuboids using a miniaturized immunoblot assay called Reverse-Phase Protein Array (RPPA) (**Fig. 6**). RPPA allowed us to analyze the state of the active kinase pathways and further validate the cuboid viability and response to drugs in hydrogel. Using cuboid dots printed on a regular well plate (**Fig. 1**A), we compared RPPA using antibodies against Phospho S-6 in Py8119 cuboids from dots analyzed with hydrogel (**Fig. 6**C), or removed from the hydrogel just before freezing for sample preparation to control for any interference of the hydrogel with processing for RPPA (**Fig. 6**B). We compared both conditions to control cuboids grown directly in medium without hydrogel (**Fig. 6**A). Phospho-S6 is a ribosomal protein and a phosphorylation marker for cell growth and protein synthesis^38^. We observed a dose-dependent decrease in phospho-S6 signal after drug treatment versus the corresponding control treatment (medium for cisplatin; 0.1 % DMSO for 2 µM staurosporine and 5 µM doxorubicin) consistent with cell death for cuboids taken from the cuboid dot and cuboids without hydrogel (**Fig. 6**A, C, D, & F). The signal from cuboids still in hydrogel seemed to be diminished compared to the cuboids alone, suggesting that the hydrogel partially interfered with the assay, particularly for the non-DMSO conditions (**Fig. 6**B&E). Replicates looked similar. Thus, our cuboid biodot approach can be used to evaluate drug treatments with not only fluorescent cell death assays, but also with the RPPA molecular protein profiling assay.

**Figure 6.**
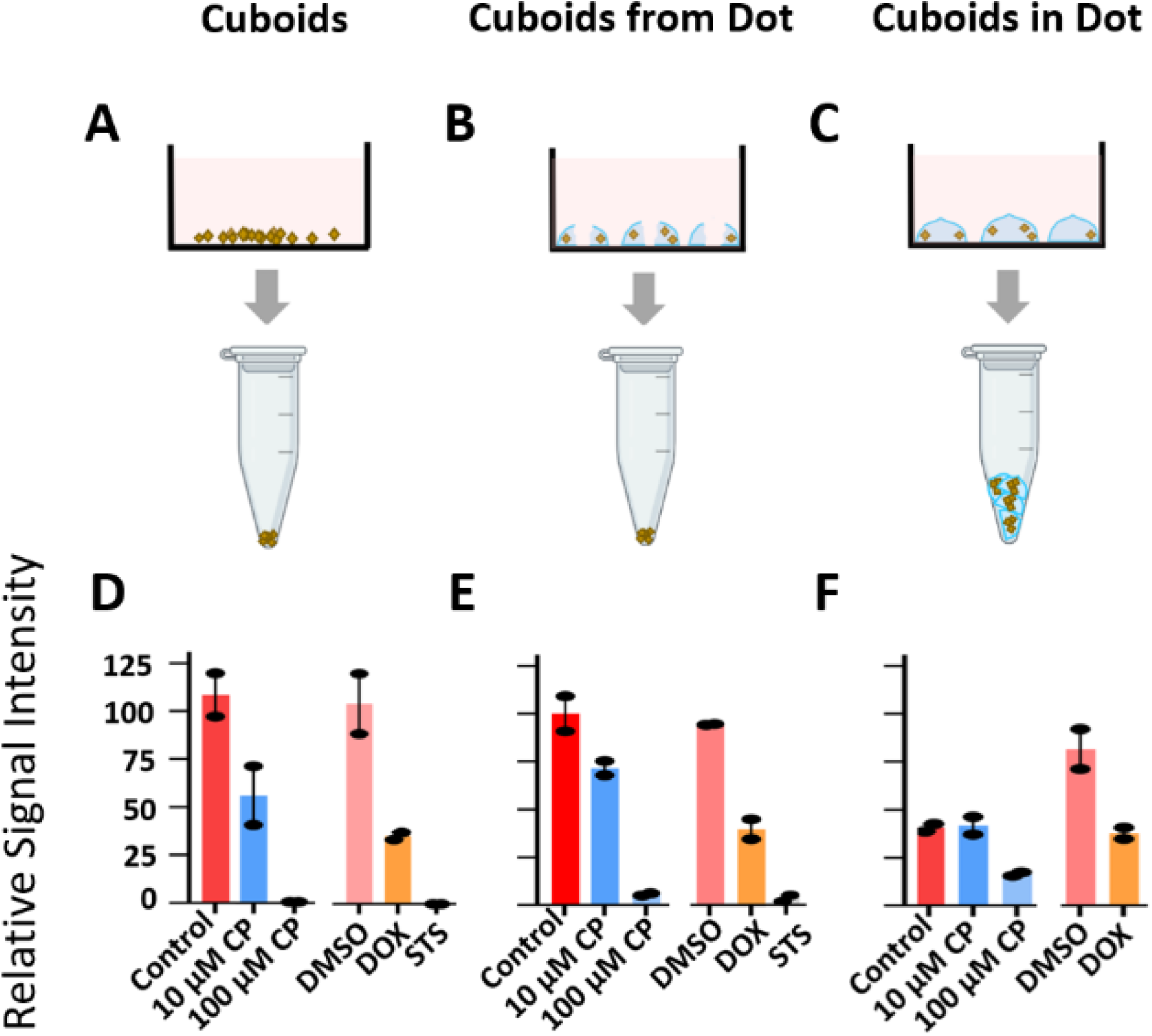
RPPA data showing phosphorylation of phospho–S6 ribosomal protein: (A-C) Reverse-phase protein array (RPPA) data showing changes in phospho-S6 after three-day drug treatments of Py8119 cuboids prepared by three different methods: control cuboids (no hydrogel), cuboids from dot (removed from hydrogel after drug exposure), and cuboids in dot (cuboids processed in hydrogel). (D-F) RPPA after 3 days drug treatment revealed changes in phospho-S6 signal, related to cell division. Replicates with ∼15 cuboids per sample.

## 4. Discussion

We have shown that microdissected tumor “cuboids” bioprinted by extrusion in hydrogel (“cuboid dots”) provide a convenient way to localize cuboids in defined locations, such as in an array. The cuboid dots can be held in place by a trap of their size or by crosslinking the hydrogel to a Transwell insert. With these simple approaches, it is possible to control the size, volume, and number of cuboids in each cuboid dot by changing straightforward bioprinting parameters such as the pressure, extrusion time, and cuboid concentration. Bioprinted cuboids in cuboid dots responded to different concentrations of cell death drugs, demonstrated through live/dead staining and RPPA.

### 4.1. CellInk ink types used for cuboid bioprinting

We considered several different ink types offered by the CellInk system, and that represent only a few of the wide range of single and multi-component hydrogels used for extrusion-based bioprinting^39^. Even though all the biomaterials (polyethylene oxide, GelMA, collagen, and alginate) that make up the bioinks tested here appear to be biocompatible with cuboids, they all should allow for rapid diffusive transport of drug through the cuboid dot, and they each have different properties^40^. We first considered a sacrificial material made of polyethylene oxide (Cellink Start). Although polyethylene oxide is a cost-effective ink, it cannot hold its shape once in aqueous form, making it suitable only to form cuboid dots for temporary placement followed by dissolution in medium. Though it is collagen-based, we did not use LifeINK, because it does not hold its form well unless printed into a gelatin slurry. As a biomolecule, collagen can regulate cell adhesion and differentiation and has been used to reconstruct bone and cartilage by bioprinting^39^. Gelatin, created by hydrolysis of collagen, offers an alternative, similarly bioactive molecule with different biophysical properties, often combined with other molecules to make a functional hydrogel^41^. GelMA is a photosensitive, gelatin-based bioink that is temperature sensitive, requiring a UV crosslinker. GelMA can encapsulate a variety of cell types to form tissue-engineered, cartilage, valves, and vessel-like structures^40,42^. It has a narrow window of optimal temperature in which it is printable, which led to too much variability in spot size for our setup. Furthermore, GelMA’s relatively high viscosity made mixing with cuboids more challenging, and the opacity of the gel made visualization of the cuboids by microscopy difficult.

For our purposes, the most suitable ink for bioprinting cuboids was a polysaccharide-derived, alginate-based bioink with cellulose nanofibrils (Cellink Bioink). We chose to use Cellink Bioink for several reasons: 1) it is easy to print and maintains its shape; 2) it crosslinks quickly; and 3) the gel is relatively cost-effective. Although cellulose primarily consists of plant-derived biomaterials, it also has excellent biocompatibility and maintains cell and cuboid viability^43^.

### 4.2. Challenges of bioprinting cuboids

We encountered a few challenges during bioprinting of cuboids in Cellink Bioink. First, the biomechanical properties of the bioink led to small variations in the dot size. The nanofabricated cellulose in Cellink enhances its shear thinning properties. Shear thinning behavior in extrusion printing improves printing by increasing flow and is beneficial for cell viability^44^. However, a time-dependent increase in flow rate due to the thixotropy of alginate (time-dependent decrease in viscosity in response to pressure)^45^ could explain the gradual increase in dot size we observed printing large numbers of dots.

Second, while mixing the ink with cuboids, air bubbles could easily become trapped in the mixture, causing less ink to extrude while bioprinting. To mitigate this problem, we gently folded the cuboids into the bioink and avoided vigorous mixing of the solution. Though CellInk’s CellMixer device, which evenly mixes cells easily and rapidly into the hydrogel, is not designed for larger microdissected tissues or organoids, its dimensions (ID 1.5 mm) are large enough for cuboids to pass through if clogging from cuboid settling could be overcome. In principle, a similar device designed for larger dimensions could be applied for use with cuboids.

Lastly, our approach of bioprinting microdissected tissues does not allow us to control the exact number of cuboids per dot. However, this challenge was not a significant limitation to our ability to bioprint cuboids for drug testing applications. Potentially one could integrate a control system as done by Mekhileri *et al*., where they used microfluidics to deposit individual large spheroids in liquid (instead of hydrogel) into preformed, bioprinted scaffolds^46^.

### 4.3. Applications of cuboid dots

The structure of cuboid dots, with microtissues in a defined hydrogel capsule, suggests many potential applications. Bioprinting allows for automated cuboid placement in precise volumes of hydrogel. Naturally, other types of microtissues could be used instead of cuboids, including non-tumor tissue (e.g., liver cuboids) or stem-cell derived organoids. Besides biocompatibility, hydrogel provides a matrix to support tissues, such as facilitation of cell to cell communication and cellular proliferation^47^. The hydrogel/tissue system of cuboid dots could be further exploited as a biological model, including addition of other cell types to the hydrogel surrounding the cuboid^48^. The presence of immune cells in cancer models and its influence on the proliferation, migration and spreading of cells is an important area to investigate^49^. In one study, researchers bioprinted melanoma cancer cells in collagen and combined them with CD8+ cells by creating microporous matrices within the hydrogel^50^. The structure facilitated immune cell spreading and a reduction in tumor cell volume^49,50^. Using tumoroids, the Le Bourhis group demonstrated infiltration of colorectal spheroids (from cell lines) by allogenic T cells and NK cells, observing spheroid apoptosis^51^. Similarly, cuboids could be interfaced with other immune cell types in a hydrogel dot to create a tumor model.

Picking up individual cuboids with tweezers or pipets can sometimes damage them and is time-consuming. Microtissues are also complex models that are highly sensitive to external factors and small modifications in protocols can lead to significant differences in results^41^. Placement of cuboids in bioprinted hydrogel dots can help automate and standardize placement of cuboids in a well plate without damage to the tissue. Furthermore, bioprinting can automate placement of microdissected tissues or organoids in well-defined spatial positions, e.g., to interface with a particular lab-on-a-chip design. The printing of larger arrays could help to facilitate high-throughput drug or biological testing with microtissues. Furthermore, cuboid dots are suitable for imaging since hydrogel is optically clear. Finally, one could also print different cuboid-encapsulating structures that interface with devices having different functionalities. For example, Melvin *et al*. developed a thiol-acrylate microfluidic resin/hydrogel hybrid system to create flow-free chemical gradients across cells^52^. The thiol-acrylate hydrogel facilitated diffusion between a sink and source channel. To generate a gradient across cuboids with the same strategy, one could replace the cell layer with a bioprinted layer of cuboids in hydrogel.

## 5. Conclusion

Using a hydrogel bioprinter, we have demonstrated that we can bioprint intact tumor cuboids in a defined spatial array, immobilized by encapsulation within hydrogel cuboid dots. We also validated the use of cuboid dots for a drug treatment assay and quantified their response using a cell death assay and RPPA. Our findings affirm the viability and drug responsiveness of cuboids in hydrogel matrices, opening the way for more advanced studies. In the future, given the suitability of hydrogels for bioengineering 3D tissues, various cell types such as immune cells and endothelial cells could be introduced into the hydrogel to analyze more complex interactions not presently possible with cuboids alone, e.g., the testing of CAR T-cell immunotherapies and/or microbe-tumor interactions.

## 6. Supporting Information Available

The following files are available free of charge. Different types of ink we used to bioprint and their properties (**Table S1**). Images of cuboids in medium, Startink, Cellink, or crosslinked Cellink and viability after Day 1 and Day 3 by Realtime Glo luminescence signal (**Fig. S1**A). Graph of viability by Day 3/ baseline ratios for each condition, where there is no significant difference between conditions (**Fig. S1**B). Graph of cell death of PY8119 mouse breast cancer cuboids either in cuboid dots (blue) or in medium (green) after 3 days of treatment with either cisplatin or medium control (**Fig. S2**).

## Author Contributions

A. B. and L. F designed, performed, and analyzed experiments and wrote the paper. T. G., C. B. L. and M. Y. performed and analyzed experiments. A. F. designed and analyzed experiments and wrote the paper.

## Institutional Review Board Statement

Not applicable.

## Informed Consent Statement

All authors of this paper are aware and agree to publish it.

## Data Availability Statement

The imaging data that support the findings of this study are available from the corresponding author upon reasonable request.

