## Supplemental Figures for "BIOPRINTING OF MICRODISSECTED TUMOR “CUBOIDS” IN HYDROGELS"

Pages: 4

Figures:2

Tables :1

**Table S1.** Types of Bioink used to Print (CelInk)

| **Table S1.** Types of Bioink used to print (CellInk) | | | | | | |
| --- | --- | --- | --- | --- | --- | --- |
| **Ink Type** | **Use** | **Solubility** | **Price** | **Crosslinker** | **Viscosity** | **Material** |
| Cellink Bioink | Used for 3-D bioprinting (tissue engineering) | Insoluble in water | Relatively expensive | Crosslinked with CaCl_2_ | Viscous (7000 Pa·s) | Alginate with hydrated cellulose nanofibrils |
| Startink | Sacrificial material | Soluble in water | Relatively inexpensive | Not crosslinkable | Not viscous (157 Pa·s) | Polyethylene oxide |
| GelMA | Use for bioprinting (cell encapsulation, tissue engineering) | Soluble in water | Relatively expensive | Crosslinked with photoinitator or exposure to UV light | Very viscous (viscosity varies with tempurature) | Porcine gelatin with methacrylate groups for crosslinking |

**Figure S1.**

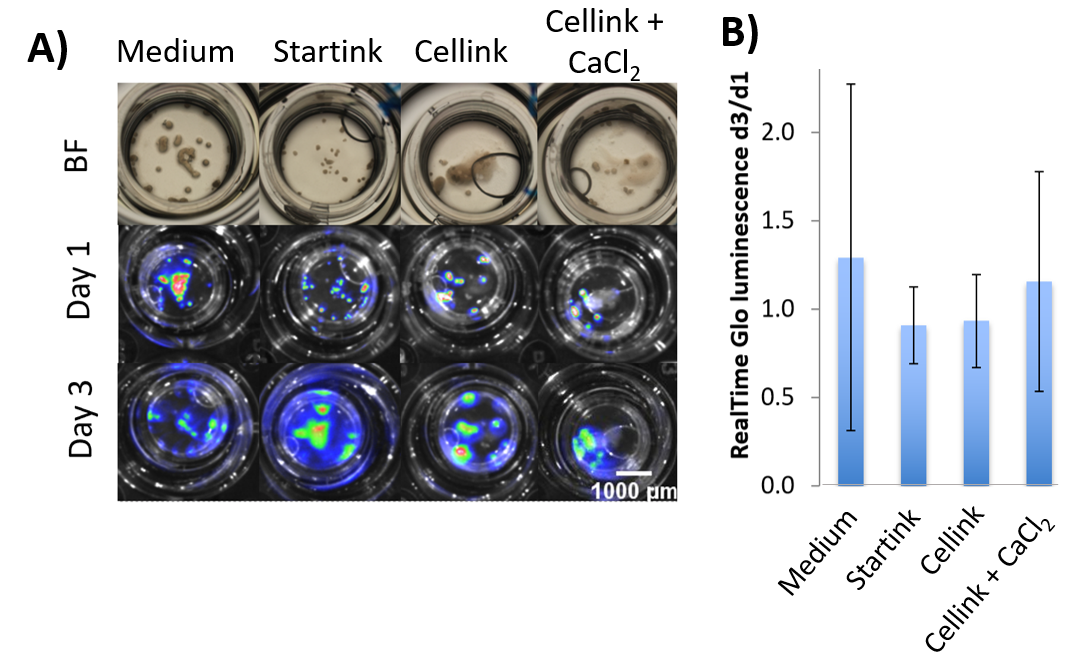

**Figure S1. Viability of cuboids in hydrogels.** (A) Images of cuboids in medium, Startink, Cellink, or crosslinked Cellink and viability after Day 1 and Day 3 by Realtime Glo luminescence signal. (B) Graph of viability by Day 3/ baseline ratios for each condition. Ave ± S.E.M. N = 12 to 32 cuboids per condition. No significant difference between conditions. One-way ANOVA with Dunnett’s multiple comparisons test.

**Figure S2.**

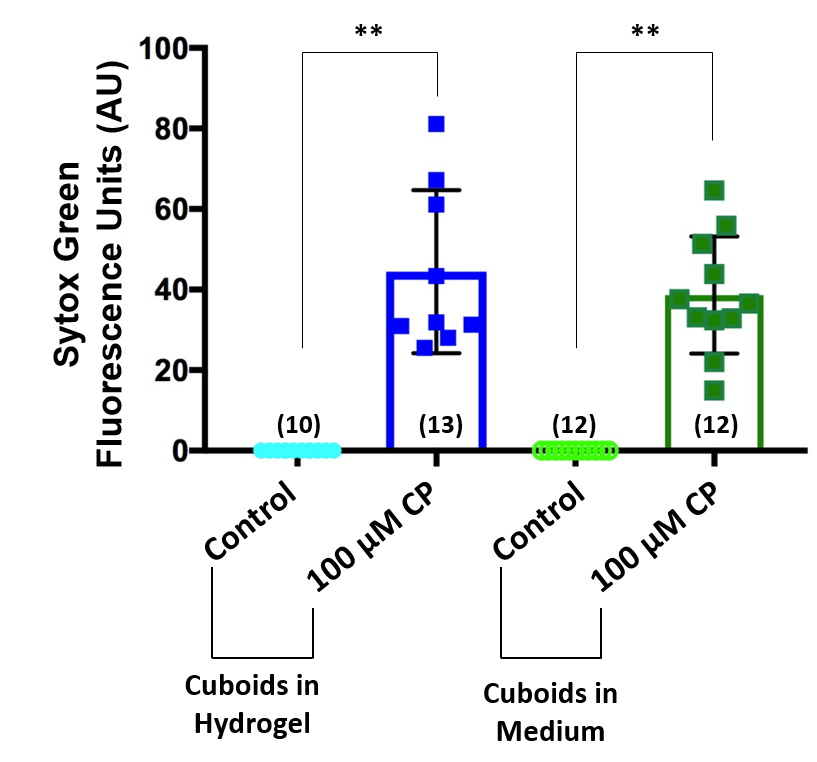

**Figure S2. Bioprinted cuboids compared to cuboids in medium.** Graph of cell death of PY8119 mouse breast cancer cuboids either in cuboid dots (blue) or in medium (green) after 3 days of treatment with either cisplatin or medium control. Mean ± St. dev was 0 ± 0 for both control conditions, 44.49 ± 19.09 cuboids in hydrogel treated with cisplatin, and 38.65 ± 13.64 for cuboids in medium without cisplatin. No significant difference between control conditions and treatment conditions. N in parenthesis. **=p< .01. One way ANOVA with Dunnett’s Multiple Comparisons test.
